## Supplementary material for "Disruption of cross-feeding interactions by invading taxa can cause invasional meltdown in microbial communities": SOM

**Supplementary Online Material:**
To investigate how model assumptions contribute to the outcome of results presented in the main text, I ran the simulations after changing two aspects of the model structure. First, I investigated how the results change when the invader is able to cross-feed. In this case, the invader is indistinguishable from native taxa, albeit with a relatively high competition value (though not higher than could be assigned by chance). Second, I allowed native taxa to have variable cross-feeding abilities, rather than assuming that metabolites were divided equally among all cross-feeders. In this case, I set cross-feeding ability as equal to competition values. For these two scenarios, I quantified the same outcomes as in the main text, and present them below.

*Sensitivity analysis 1: Invaders have the same cross-feeding abilities as native taxa*

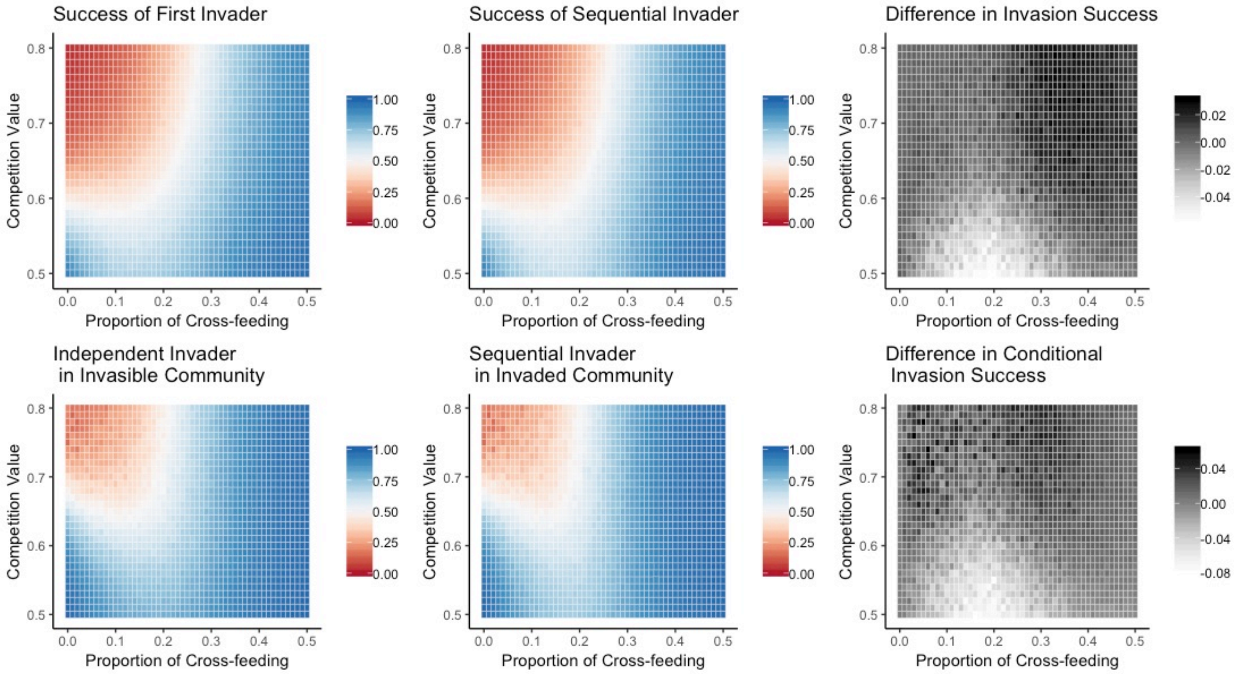


**Fig. S1: Same results as presented in Fig. 3 in the main text, but under the condition that invaders have equivalent cross-feeding dynamics as native taxa.**

Allowing invasive taxa to cross-feed substantially changes invasion success. When invaders can cross-feed, primary invasions are much more likely to be successful (upper left panel); in fact, higher rates of cross-feeding in the community facilitate the invader. Secondary invaders are, overall, slightly less successful than primary invaders (upper middle panel), and the greatest discrepancy between primary and secondary invaders occurs at intermediate levels of cross-feeding (upper right panel). This result that a primary invasion can make a later invasion more difficult is in direct contrast to the results presented in the main text. Thus, the cross-feeding dynamics of an introduced taxon has strong influence on both its ability to join the community and the potential for further taxa to join the community.

When looking only at communities that were successfully invaded, a different invader is generally also successful there (lower left panel). A secondary invader shows similar success patterns to a primary invader (lower middle panel), but there are regions of parameter space where the secondary invader has both greater and lesser success (lower right panel). In general, when competition is low, secondary invaders were less successful, especially at intermediate levels of crossfeeding. However, when competition was high, a secondary invader was often more successful, due to the presence of the primary invader.

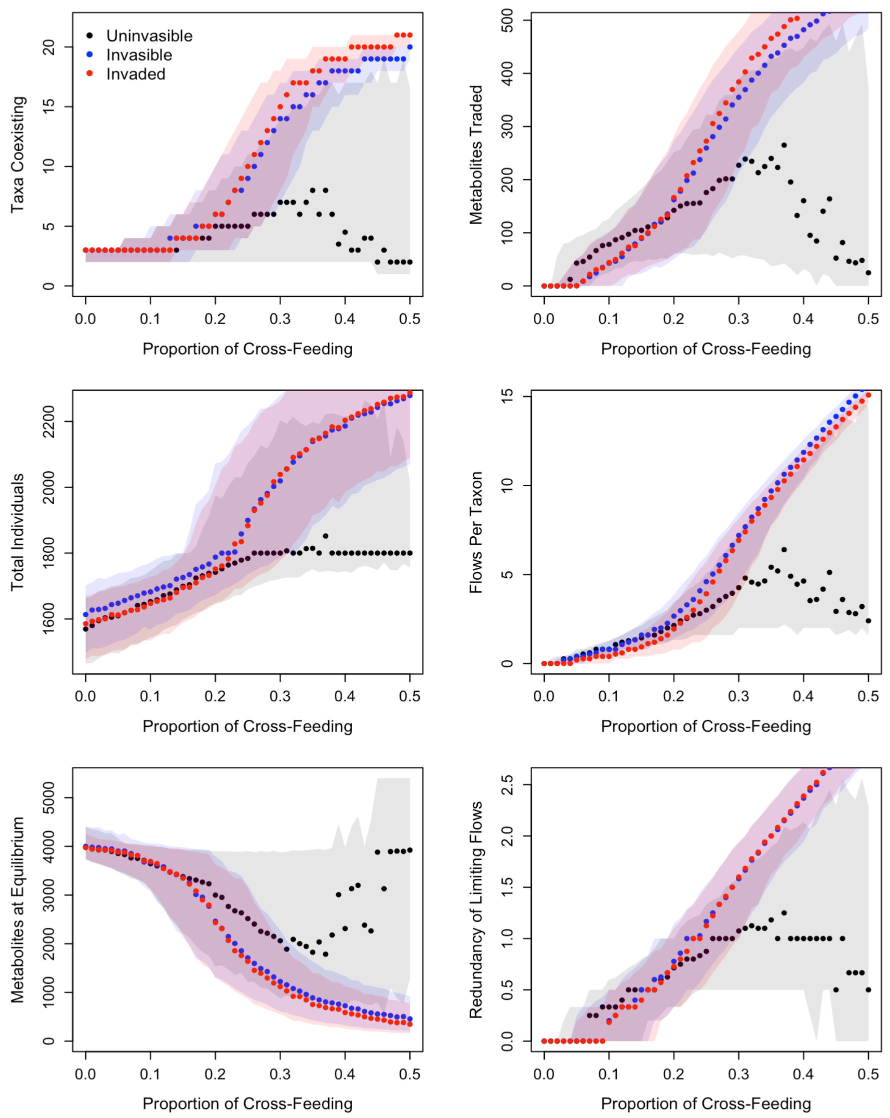


**Fig. S2: Same results as presented in Fig. 5 in the main text. When invaders are able to cross-feed, a successful invasion only minimally changes community structure and metabolite exchange networks.**

In contrast to Fig. 5 in the main text, which showed that the presence of an invader changes both community structure and metabolite exchange dynamics, Fig. S2 shows that the introduction of an invader generally has small effects on the communities. When native communities are already highly diverse (i.e. have a large number of taxa coexisting), the invader often simply joins the community, without dislodging other taxa (upper left panel). In these cases, the number of metabolites traded increases (upper right panel), as another taxon is added to the network of metabolite transfers.

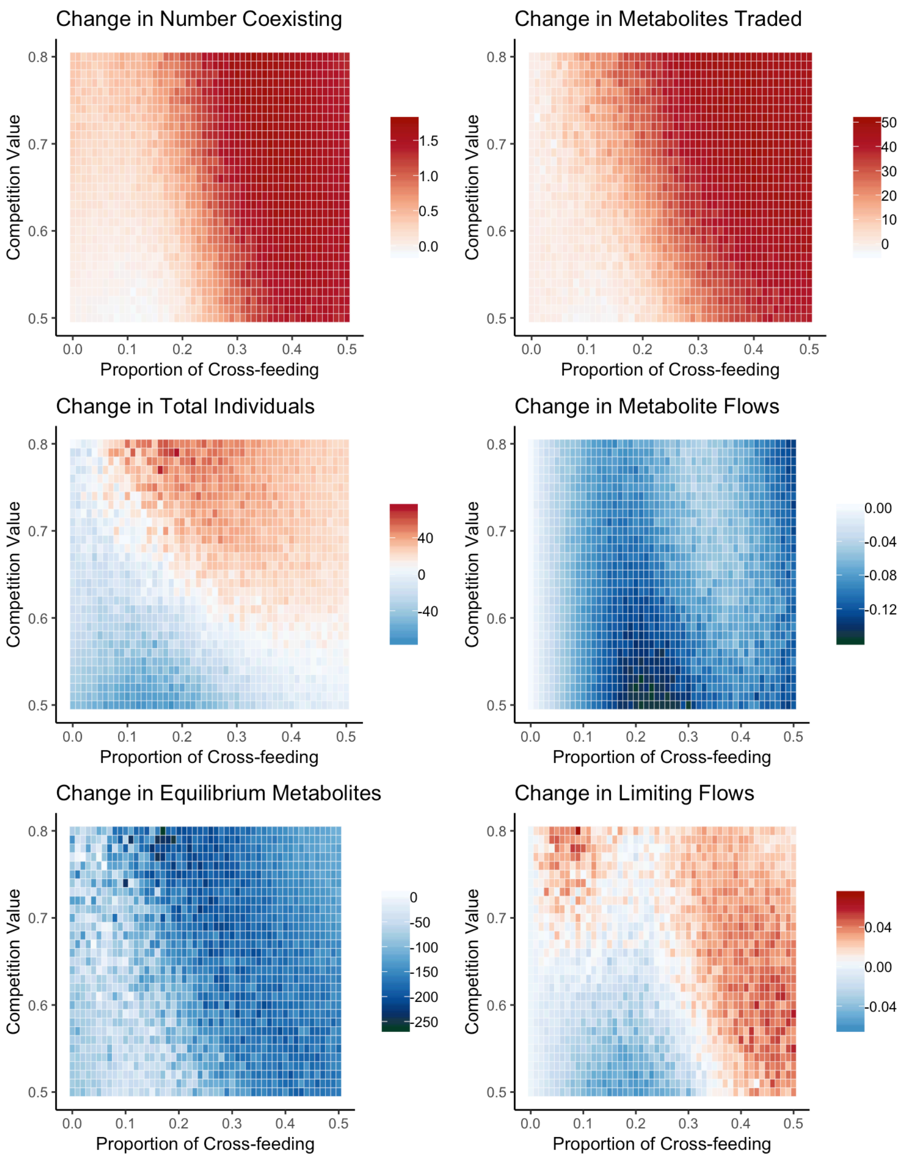


**Fig. S3: Same results as presented in Fig. 6 in the main text. Warm colors indicate increases, and cool colors indicate decreases.**

Allowing the invaders to cross-feed alters the impacts of a successful invasion on community structure and metabolite exchange. When invaders can cross-feed, they are much less likely to remove taxa from communities or diminish the number of metabolites exchanged within the community (upper panels). Additionally, there is less change in the total number of taxa in the community (left middle panel). Invading taxa also generally lead to decreases in the number of equilibrium metabolites (bottom left panel). Finally, invaders have a smaller impact on overall cross-feeding networks, although they do generally lead to slightly fewer cross-feeding relationships overall (right middle panel). However, the effect of invaders on the number of metabolite flows providing limiting nutrients could be either positive or negative, depending on the combination of cross-feeding and competition parameters (bottom right panel).

*Sensitivity Analysis 2: Cross-feeding ability is equal to competitive ability*

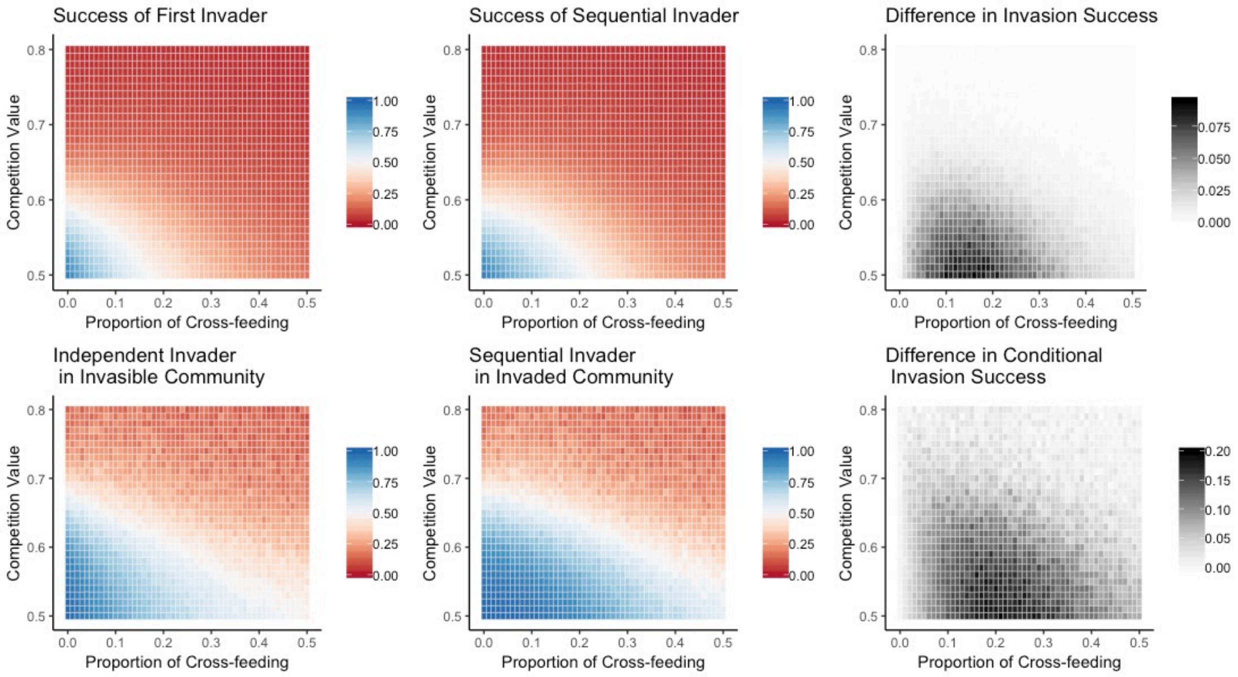


**Fig. S4: Same results as presented in Fig. 3 in the main text, but under the condition that cross-feeding abilities are variable between taxa, being set as equal to each taxon's competition value.**

In these simulations, the distinction from the model presented in the main text is that taxa in the native community have varying levels of cross-feeding abilities. However, this change has minimal impact on invasive taxa, as invasion success is largely unchanged.

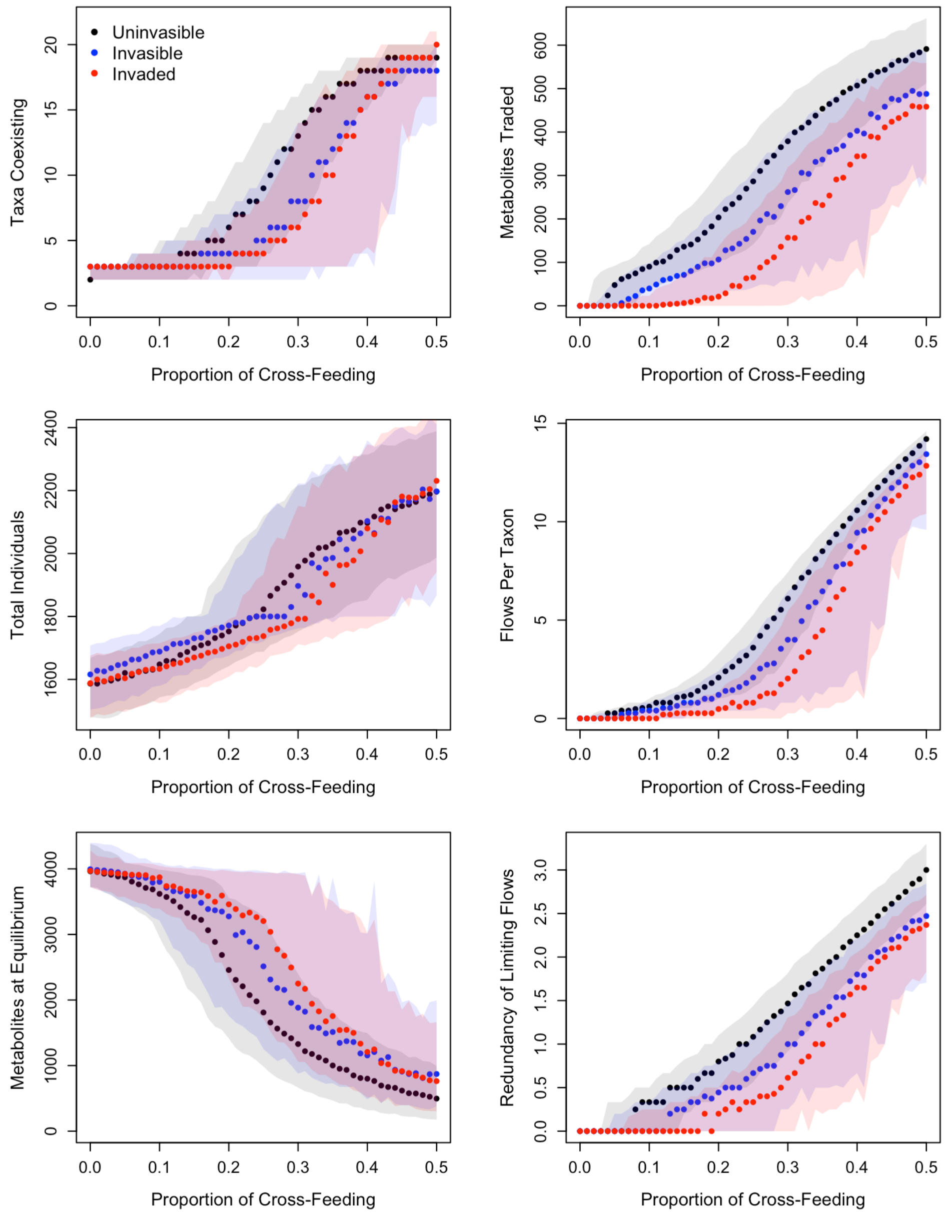


**Fig. S5: Same results as presented in Fig. 5 in the main text. Allowing for variation of cross-feeding abilities in the native taxa has minimal effect on model results.**

As with invasion success, changing the model to allow native taxa to be differentially good at cross-feeding shows minimal differences in community structure and metabolite transfer networks.

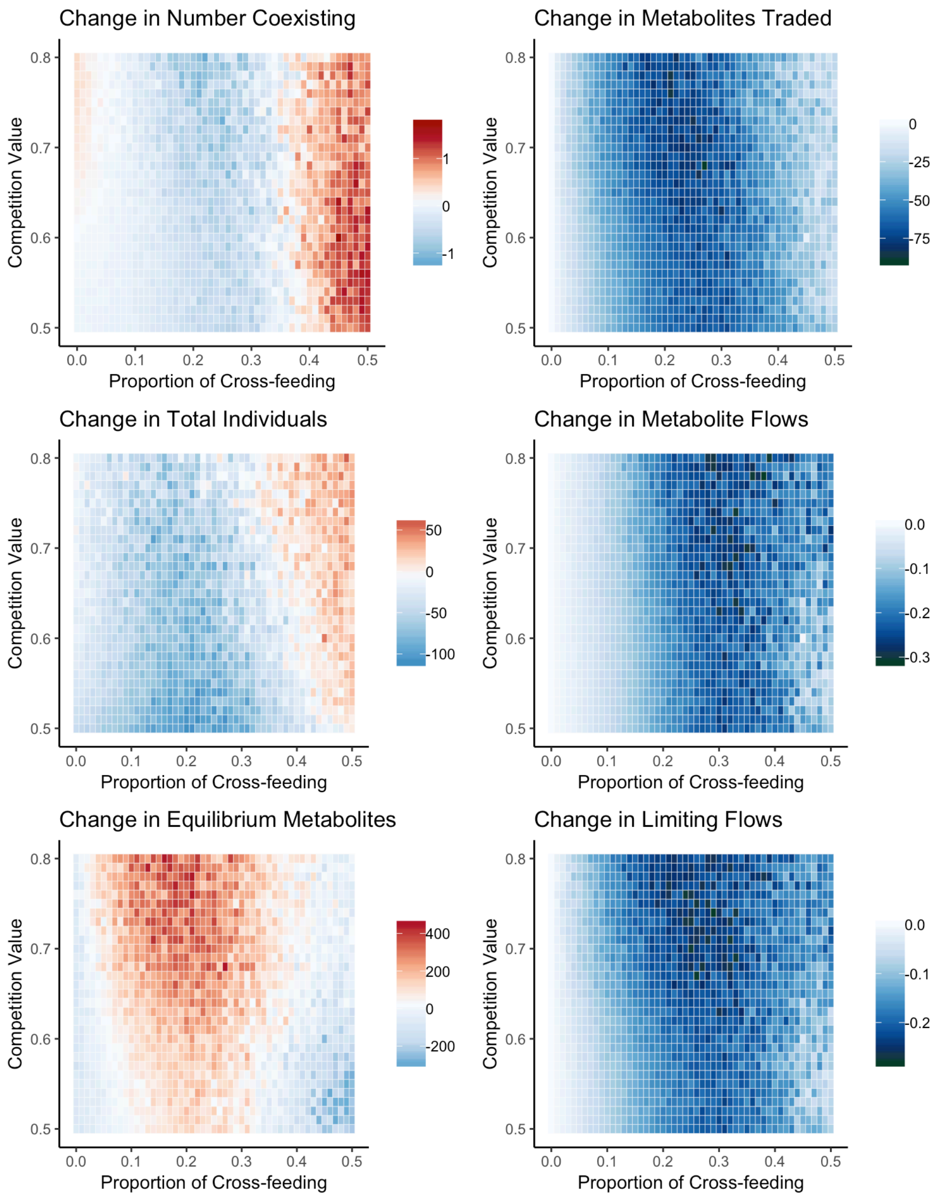


**Fig. S6: Same results as presented in Fig. 6 in the main text.**

Finally, Fig. S6 looks at how communities changed in response to a successful invader, when the native taxa varied in cross-feeding ability. These results are, again, largely similar to those presented in the main text, where there is no variation in cross-feeding ability.
